# Cerebral contribution to the execution, but not recalibration, of motor commands in a novel walking environment

**DOI:** 10.1101/686980

**Authors:** D. de Kam, P.A. Iturralde, G. Torres-Oviedo

## Abstract

Human movements are flexible as they continuously adapt to changes in the environment by updating planned actions and generating corrective movements. Planned actions are updated upon repeated exposure to predictable changes in the environment, whereas corrective responses serve to overcome unexpected environmental transitions. It has been shown that corrective muscle responses are tuned through sensorimotor adaptation induced by persistent exposure to a novel situation. Here, we asked whether cerebral structures contribute to this recalibration using stroke as a disease model. To this end, we characterized changes in muscle activity in stroke survivors and unimpaired individuals before, during, and after walking on a split-belt treadmill moving the legs at different speeds, which has been shown to induce recalibration of corrective responses in walking in healthy individuals. We found that the recalibration of corrective muscle activity was comparable between stroke survivors and controls, which was surprising given then known deficits in feedback responses post-stroke. Also, the intact recalibration in the group of stroke survivors contrasted the patients’ limited ability to adjust their muscle activity during steady state split-belt walking compared to controls. Our results suggest that the recalibration and execution of motor commands in new environments are partially dissociable: cerebral lesions interfere with the execution, but not the recalibration, of motor commands upon novel movement demands.

## Introduction

Humans continuously adapt their movements to changes in the body or environment through corrective responses and adjustment of planned actions. Corrective responses are rapidly triggered upon unexpected movement disturbances (Jordan and Rumelhart, 1992; Bhushan and Shadmehr, 1999). Conversely, planned actions are predictive in nature and are updated through sustained perturbations (e.g., constant force) altering one’s movement (Wolpert *et al.*, 1998). Recent work has shown that corrective motor commands also adapt to persistent changes in the environment, such that they become appropriate to the novel situation at hand (e.g. Iturralde and Torres-Oviedo, 2019). However, little is known about the neural processes contributing to the recalibration of corrective responses.

It has been suggested that planned and corrective actions share an internal representation of the environmental dynamics (Wagner and Smith, 2008; Maeda *et al.*, 2018), thus their recalibration could rely on updates to these internal models (Wolpert *et al.*, 1998). If so, the recalibration of corrective responses is likely dependent on cerebellar (Smith and Shadmehr, 2005; Morton and Bastian, 2006), but not cerebral structures (Reisman *et al.*, 2007; Choi *et al.*, 2009). However, tuning of corrective responses according to the environmental dynamics is cerebral-dependent (Trumbower *et al.*, 2013; de Kam *et al.*, 2018) and several studies have shown that corrective muscle responses are affected post-stroke (Marigold and Eng, 2006; De Kam *et al.*, 2017; de Kam *et al.*, 2018). Thus, it is plausible that the recalibration of corrective responses is also affected after cerebral lesions, which would imply that cerebral structures contribute to the adaptation of corrective actions. Here, we evaluate the involvement of cerebral structures in the recalibration of reactive control through the analysis of corrective muscle activity in individuals with cerebral-lesions after stroke.

We characterized stroke-related deficits in muscle activity before, during, and after split-belt walking, which induces robust locomotor adaptation (Reisman *et al.*, 2007). We hypothesized that the execution of motor patterns in a novel walking situation and the subsequent recalibration of corrective responses would be impaired post-stroke based on literature indicating that muscle patterns are generally affected after cerebral lesions (Bowden *et al.*, 2010; Clark *et al.*, 2010; Cheung *et al.*, 2012; de Kam *et al.*, 2018). This would suggest that cerebral structures are involved in both the execution of motor commands in a novel situation and the recalibration of corrective actions that results from extended exposure to the novel environmental demands.

## Methods

### Subjects

We tested 16 stroke survivors in the chronic phase (> 6 months) with unilateral supratentorial lesions (i.e. without brainstem or cerebellar lesions; Age 62±9.9 years, 6 Females, Table 1) and 16 age and gender matched controls (Age 61±9.7 years, 6 Females). We applied the following inclusion criteria: 1) be able to walk with or without a hand-held device at a self-paced speed for at least 5 minutes, 2) have no orthopedic or pain conditions interfering with the assessment, 3) have no neurological conditions except stroke, 4) have no severe cognitive impairments (defined as mini-mental state exam < 24), 5) have no contraindications for performing moderate intensity exercise and 6) use no medication that interferes with cognitive function. We excluded from data analysis 4 out of the 32 participants invited for testing. One stroke participant (P7) was excluded because of severe muscle atrophy and weakness on the sound limb (i.e., non-paretic side), which was present prior to the brain lesion. Another stroke participant (P3) was excluded because of poor muscle recordings due to technical difficulties during testing. One control participant (C1) was excluded because this person failed to follow the testing instructions. Lastly, we had to remove C7 (i.e. age-matched control of P7) because our regression analyses required equal sample sizes across groups. Namely, including fewer participants in the regression of one group reduces the regressor estimates due to more noise in the averaged data. The study protocol was approved by the Institutional Review Board at the University of Pittsburgh. All study participants gave written informed consent prior to participation.

**Table 1.**
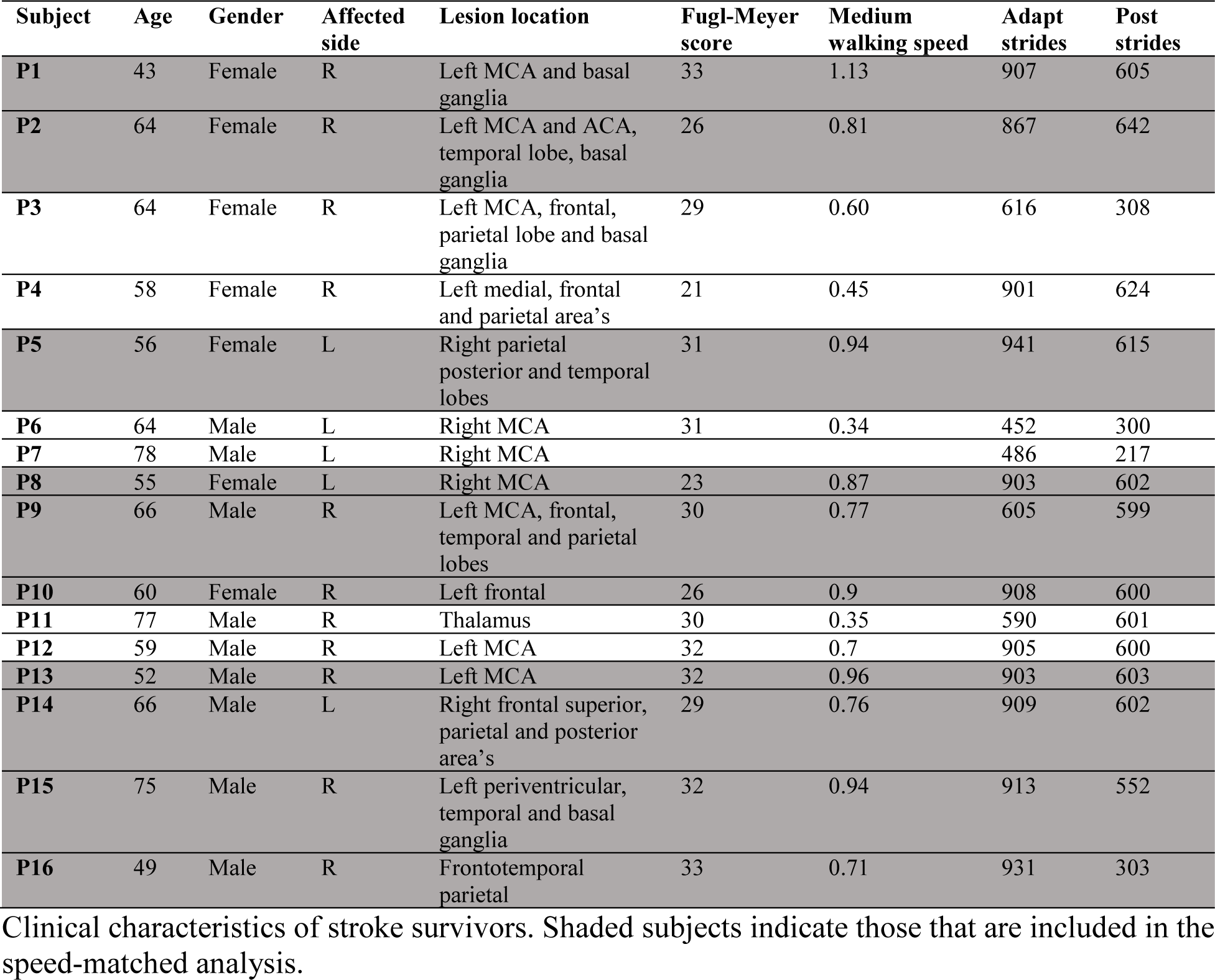
Clinical characteristics of stroke survivors.

| Subject | Age | Gender | Affected side | Lesion location | Fugl-Meyer score | Medium walking speed | Adapt strides | Post strides |
| --- | --- | --- | --- | --- | --- | --- | --- | --- |
| **P1** | 43 | Female | R | Left MCA and basal ganglia | 33 | 1.13 | 907 | 605 |
| **P2** | 64 | Female | R | Left MCA and ACA, temporal lobe, basal ganglia | 26 | 0.81 | 867 | 642 |
| **P3** | 64 | Female | R | Left MCA, frontal, parietal lobe and basal ganglia | 29 | 0.60 | 616 | 308 |
| **P4** | 58 | Female | R | Left medial, frontal and parietal area's | 21 | 0.45 | 901 | 624 |
| **P5** | 56 | Female | L | Right parietal posterior and temporal lobes | 31 | 0.94 | 941 | 615 |
| **P6** | 64 | Male | L | Right MCA | 31 | 0.34 | 452 | 300 |
| **P7** | 78 | Male | L | Right MCA |  |  | 486 | 217 |
| **P8** | 55 | Female | L | Right MCA | 23 | 0.87 | 903 | 602 |
| **P9** | 66 | Male | R | Left MCA, frontal, temporal and parietal lobes | 30 | 0.77 | 605 | 599 |
| **P10** | 60 | Female | R | Left frontal | 26 | 0.9 | 908 | 600 |
| **P11** | 77 | Male | R | Thalamus | 30 | 0.35 | 590 | 601 |
| **P12** | 59 | Male | R | Left MCA | 32 | 0.7 | 905 | 600 |
| **P13** | 52 | Male | R | Left MCA | 32 | 0.96 | 903 | 603 |
| **P14** | 66 | Male | L | Right frontal superior, parietal and posterior area's | 29 | 0.76 | 909 | 602 |
| **P15** | 75 | Male | R | Left periventricular, temporal and basal ganglia | 32 | 0.94 | 913 | 552 |
| **P16** | 49 | Male | R | Frontotemporal parietal | 33 | 0.71 | 931 | 303 |

Clinical characteristics of stroke survivors. Shaded subjects indicate those that are included in the speed-matched analysis.

### Experimental setup and protocol

We investigated how participants adapted their kinematic and muscle activation patterns on an instrumented split-belt treadmill (Bertec, Columbus, Ohio, USA) with two belts that moved at either the same speed (tied condition) or at different speeds (split condition). We kept the mean speed across the belts constant in the tied and split conditions. Each subjects’ mean belt speed was set to 0.35 m/s below their overground walking speed during the 6 minutes walking test (Rikli and Jones, 1998; Kervio *et al.*, 2003), yielding a comfortable speed for treadmill walking. The mean belt speed, denoted as medium speed, is reported for each subject in Table 1. In the split condition, the speed of one belt was decreased (slow belt) and the speed on the other belt was increased (fast belt) by 33% of the medium speed to obtain a belt speed ratio of 2:1. Stroke survivors walked with their paretic leg on the slow belt, whereas healthy subjects walked with their non-dominant leg on the slow belt. The treadmill protocol consisted of 5 periods: 1) 50 strides (e.g. time between two subsequent heel strikes of the same leg) walking at medium speed, 2) a Short Exposure (10 strides) to the split condition, 3) 150 strides of Baseline walking at medium speed, 4) 900 strides of Adaptation to the split condition and 5) 600 strides of Post-Adaptation at medium speed (Figure 1A). Subjects had several resting breaks during the experiment and some stroke individuals completed fewer strides during Adaptation and Post-Adaptation to prevent fatigue (Table 1 shows number of strides completed per subject). Participants wore a safety harness, not supporting body-weight, attached to a sliding rail in the ceiling to prevent falls. Moreover, subjects could hold on to a handrail in front of the treadmill, but were instructed to do so only if needed.

**Figure 1.**
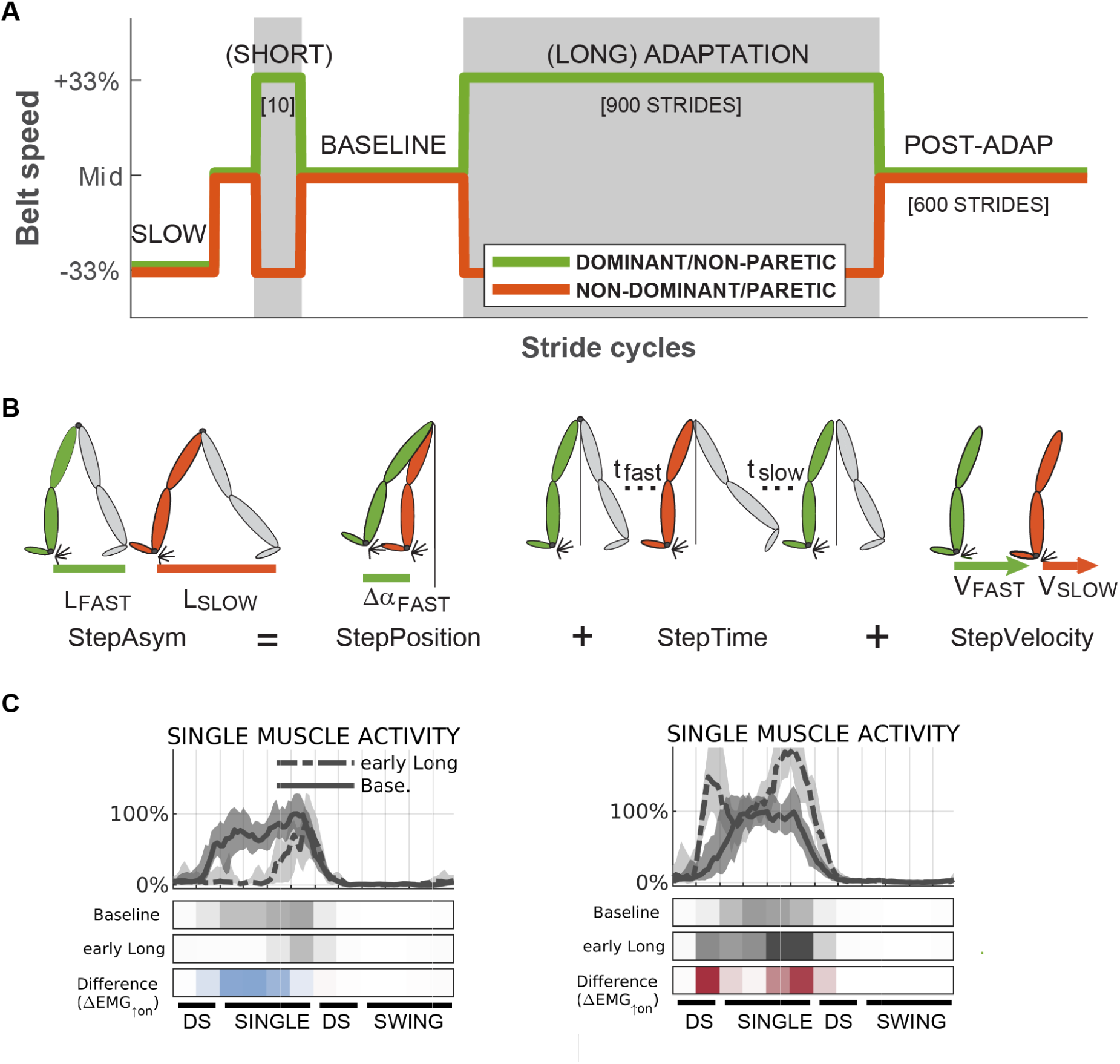
Overview of experimental methods. **A)** Schedule of belt speeds experienced by subjects. **B)** Schematic representation of definitions of kinematic parameters StepAsym, StepPosition, StepTime and StepVelocity, adapted from (Sombric *et al.*, 2017) **C)** Sample EMG traces of one muscle (LG) during Baseline and late Adaptation for a representative control subject. Median activity across strides (lines), and the 16-84 percentile range (shaded). Data was lowpass filtered for visualization purposes. Colorbars below the traces represent averaged normalized values during 12 kinematically-aligned phases of the gait cycle (see Methods) for Baseline, early Adaptation, and the difference (red indicates increase, blue decrease).

### Data collection

We collected kinetic, kinematic, and electromyography (EMG) data to characterize individuals’ walking pattern. The ground reaction force aligned with gravity (Fz, sampled at 1000Hz) was used to identify the instants at which the feet landed (i.e., heel-strike: Fz>10N) or were lifted from the ground (i.e. toe-off: Fz<10N) (Iturralde and Torres-Oviedo, 2019). The positions of the ankles (lateral malleolus) and hips (greater trochanter) were recorded at 100Hz using a 3D motion analysis system (Vicon Motion Systems, Oxford, UK). Activity of 15 muscles (See Supplementary Table 1) was recorded bilaterally at 2000Hz using a Delsys Trigno System (Delsys Inc., Natick, Massachusetts). EMG signals were high-pass filtered with a 30Hz 4^nd^ order Butterworth dual-pass filter and subsequently rectified (Merletti and Parker, 2005).

### Data analysis

#### Kinematic parameters

We characterized the adaptation of step length asymmetry (StepAsym, Eq1, Figure 1B), which is conventionally used to quantify gait changes during split-belt walking (Reisman *et al.*, 2007; Torres-Oviedo *et al.*, 2011). We defined StepAsym as the difference between consecutive steps of the legs in terms of step length, where step length is the distance between the feet (i.e., ankle markers) at heel strike. In our definition, StepAsym is positive when the step length of the fast leg (i.e. dominant or non-paretic) is larger than the one of the slow leg (non-dominant or paretic). We also quantified spatial (StepPosition) and temporal (StepTime) gait features that contribute to StepAsym, since those are differentially affected across stroke survivors and they exhibit distinct adaptation patterns in unimpaired adults during split-belt walking (Finley *et al.*, 2015). Finally, StepVelocity was defined as the difference between the legs in terms of velocity of the foot with respect to the body when in contact with the ground. All parameters were expressed in units of distance and they were normalized to the sum of left and right step lengths in order to account for differences in step sizes across subjects (Sombric *et al.*, 2017).

#### EMG parameters

We characterized the modulation of muscle activity across the different walking conditions using the average activity of each muscle for fixed phases of the gait cycle (Figure 1C). Specifically, we divided the gait cycle into 4 phases: first double support (DS; between ipsilateral heel strike and contralateral toe off), single stance (SINGLE; from contralateral toe-off to contralateral heel-strike), second double support (DS; between contralateral heel strike and ipsilateral toe off) and swing (SWING; between ipsilateral toe-off and ipsilateral heel-strike). We further divided each of these phases to achieve better temporal resolution. Specifically, both DS phases were divided in two equal sub-phases and the SINGLE and SWING phases were sub-divided in four equal sub-phases. Muscle activity amplitude was averaged in time for each of these subintervals for every stride and muscle resulting in 180 muscle activity variables per leg per stride cycle: 12 subinterval × 15 muscles.

EMG activity for each muscle was linearly scaled to baseline walking (last 40 strides), such that a value of 0 corresponded to the average of the interval with the lowest average activity and 1 corresponded to the average of the interval with the highest average activity (Iturralde and Torres-Oviedo, 2019). This normalization enabled us to aggregate the EMG activity across subjects to perform group analyses. Of note, we excluded from analysis the activity of soleus from one stroke survivor because technical difficulties during data collection.

#### Epochs of interest

Kinematic and EMG parameters were used to characterize subjects’ behavior at the beginning (‘early’) and at the end (‘late’) of each experimental condition. Specifically, the epochs of interest included: late Baseline walking, early and late Adaptation and early Post-adaptation. The ‘early’ epochs were characterized by the median of the initial 5 strides and ‘late’ epochs by the median of the last 40 strides of the condition of interest. We chose medians across strides, rather than means to minimize the impact of outlier values. In all cases, we excluded the very first and very last stride of each condition to avoid artifacts from starting and stopping the treadmill. Subsequently, we subtracted the late Baseline behavior from all epochs of interest. This allowed us to identify group differences in subjects’ modulation of kinematic and EMG parameters beyond those due to distinct baseline biases. Moreover, we computed the differences between EMG activity early Post-Adaptation vs. late Adaptation to quantify changes in EMG activity upon sudden removal of the perturbation.

#### Sensorimotor recalibration of corrective muscle responses

We studied the structure (i.e., activity across multiple muscles) of corrective motor responses upon sudden changes in the walking environment (Figure 2), since this reflects the extent of sensorimotor recalibration (Iturralde and Torres-Oviedo, 2019). We defined corrective responses as the rapid changes in motor output (ΔEMG) immediately after a transition in the walking environment. Corrective responses were quantified as the difference in muscle activity immediately after an environmental transition (EMG_after_) compared to the muscle activity before the transition (EMG_before_, Figure 2A). Thus, corrective response (ΔEMG)=EMG_after_–EMG_before_. Since we had multiple strides before and after a transition, we used the median EMG activity across either 40 or 5 strides to quantify EMG_before_ and EMG_after_ a given transition, respectively. For example, ΔEMG_on(+)_ indicated the corrective response upon introducing a split-belt environment in which the dominant (or non-paretic) leg moved faster than the non-dominant (or paretic) leg, which was an environment arbitrarily defined ‘(+)’ (Figure 2A). Thus, ΔEMG_on(+)_ was computed as the difference between EMG activity before and after the ‘+’ environment was introduced.

**Figure 2.**
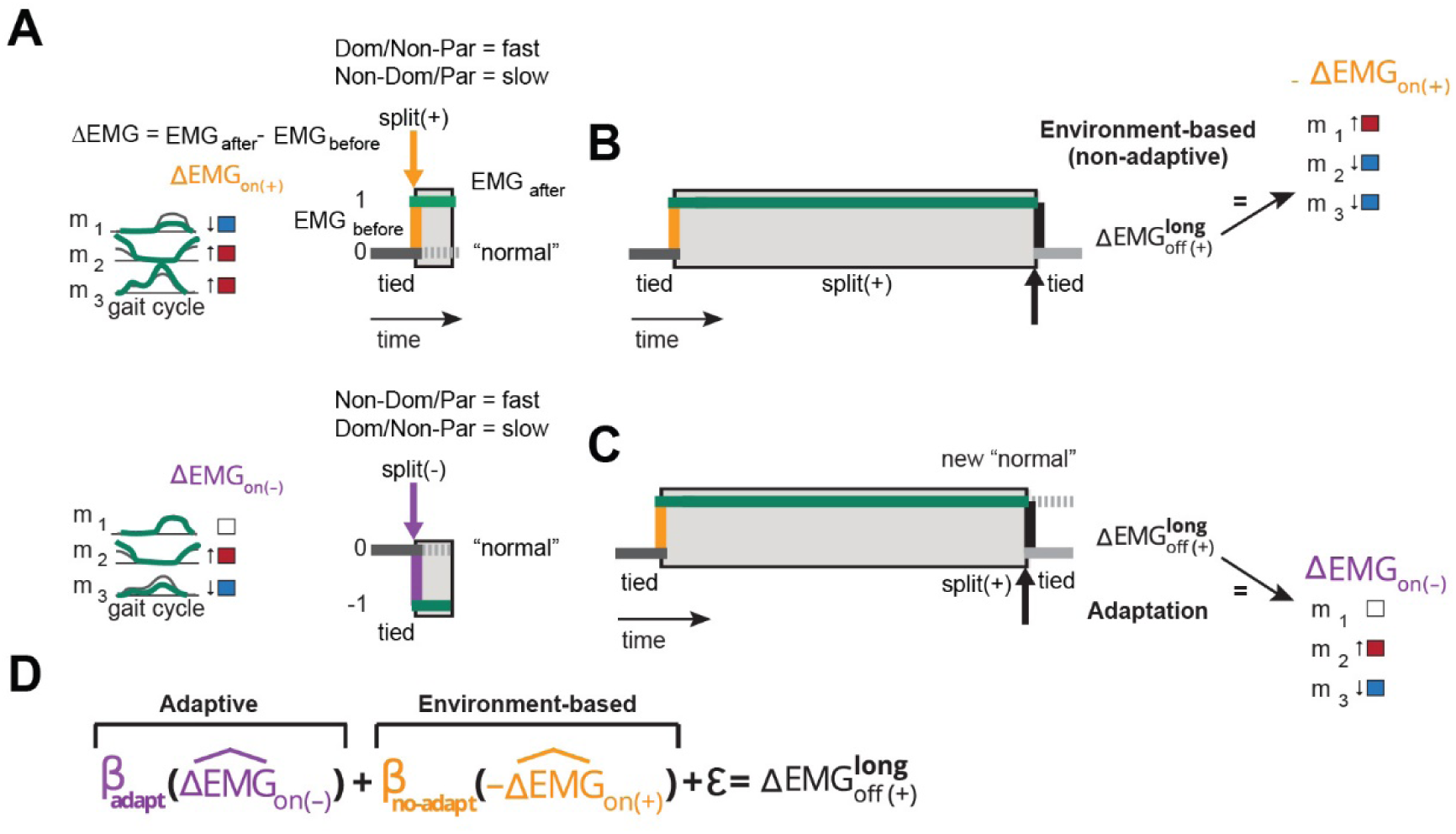
Environment-based and adaptive contributions to corrective responses. **A)** Schematic representation of corrective responses upon the introduction split-belt perturbation. The split environment is arbitrarily defined as “+” if the paretic leg is on the slow belt and the non-paretic one is on the fast belt (upper cartoon), whereas it is defined as “-” if the paretic leg is on the fast belt and the non-paretic one is on the slow belt. Changes in muscle activity (i.e. corrective response) upon the introduction of the “+” or “-” environment are color-coded as blue (decreased activity), red (increased activity) and white (no change in activity). **B)** In the case of an environment-based corrective response changes in muscle activity perturbation removal (ΔEMG_off(+)_) are opposite to those upon perturbation introduction. **C)** In the case of an adaptive corrective response, the split-environment is perceived as the new normal. Consequently, removal of the split-environment will be experienced as a perturbation in the opposite direction. Thus, the structure of the corrective response will resemble the one observed upon introduction of the “-” environment. **D)** Regression equation used to quantify the structure of corrective response ΔEMG_off(+)_. In this equation, β_adapt_ quantifies the similarity of ΔEMG_off(+)_ to the adaptation-based response and β_no-adapt_ quantifies the similarity of ΔEMG_off(+)_ to the environment-based response.

We were specifically interested in the structure of corrective responses post-adaptation because this structure indicates the extent to which subjects recalibrate their motor system (Iturralde and Torres-Oviedo, 2019). Namely, the structure of these corrective responses is determined by both changes in the environment and changes in the motor systems’ adaptive state. We discerned the environment-based and adaptive-based contributions to corrective responses post-adaptation (ΔEMG_off(+)_) with a regression model (ΔEMG_off(+)_ = adaptive-based + environment-based + ε). In the case of an environment-based response, the corrective pattern ΔEMG_on(+)_ upon introducing the ‘+’ split environment is simply disengaged once this environment is removed (i.e., both belts moving at the same speed (Iturralde and Torres-Oviedo, 2019)). Thus, in this case the structure of corrective responses post-adaptation ΔEMG_off(+)_ (i.e., when the split ‘+’ environment is turned off) resembles the numerical opposite of ΔEMG_on(+)_ (ΔEMG_off(+)_=-ΔEMG_on(+)_ Figure 2B). Conversely, adaptive-based responses are observed if subjects perceive the novel environment ‘+’ as the ‘new normal’, such that removing it is equivalent to transitioning into a split condition in the opposite direction (i.e., non-dominant leg moving faster than the dominant one, Figure 2A). Thus, in the case of adaptive corrective responses, the structure of ΔEMG_off(+)_ resembles corrective responses to transitioning into the opposite ‘-’ split-belt environment in which the paretic or non-dominant leg would increase speed and the non-paretic or dominant one decreases it (ΔEMG_off(+)_=ΔEMG_on(-)_; Figure 2C). We concurrently quantified the degree of sensorimotor recalibration for each leg with the following regression equation (Figure 2D):

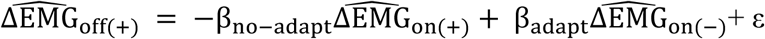

In the regression equation, the parameters β_no-adapt_ and β_adapt_ are respectively interpreted as the extent to which the structure of corrective responses indicates transitions in the environment (i.e., environment-based) or the adaptation of subjects’ motor system (i.e., adaptive-based). Note that every vector is divided by its norm (i.e.,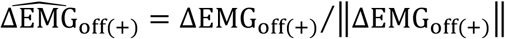). This was done because we were interested in identifying stroke-related deficits in the structure, rather than the magnitude of corrective responses, which is known to be different (e.g. De Kam *et al.*, 2017). For example, we find that the amplitude of corrective responses ΔEMG_on(+)_ for each leg was smaller for the stroke (∥ΔEMG_on(+)_∥=2.6 and 2.2) than the control group (3.3 and 3.7).

Note that ΔEMG_on(-)_ was not directly measured to avoid exposing subjects to multiple environmental transitions prior to the Adaptation period. Instead, we inferred these responses by exploiting the symmetry of the transition between the two legs. The only difference between the ‘+’ and ‘-’ environments is which leg increases speed and which leg decreases it. We used this similarity to infer the (not recorded) corrective responses (ΔEMG_on(-)_) of each leg to transitioning into the ‘-’ environment from the (measured) corrective responses (ΔEMG_on(+)_) to transitioning to the ‘+’ environment. In other words, we assumed that the (not recorded) non-dominant leg’s responses to the “on (-)” transition would be similar to the (recorded) dominant leg’s responses to the “on (+)” transition, and vice versa. We are aware that this assumption might not be valid for some post-stroke individuals, given their inherent motor asymmetry. Thus, group differences in β_adapt_ values, which are estimated using the not recorded ΔEMG_on(-)_ in our regression analysis, might be due to the experimental limitation of our study. To address this possibility, we performed a post-hoc analysis to compare the regression coefficients between a subset of patients and controls (n=7 on each subgroup) that had similar asymmetry in their EMG activity during baseline walking (p=0.1).

#### Structure of muscle activity patterns in a novel walking environment

We characterized changes in the structure of steady state muscle activity from baseline walking to late Adaptation (ΔEMG_SS_ = EMG_late Adaptation_ – EMG_late Baseline_). This was defined as the pattern of activity across all muscles and all gait cycle intervals (15 muscles × 12 intervals = 180 data points for a given epoch). The ΔEMG_SS_ 180-dimensional vector for each subject was used to assess structural differences between stroke survivors and controls. We specifically computed a cosine between the ΔEMG_SS_ for each individual and a ‘reference pattern’ ΔEMG_SS_, which was defined as the median ΔEMG_SS_ of the control group. This reference pattern for ΔEMG_SS_ was calculated as the group median of all control subjects when computing the similarity metric for each leg of the stroke survivors, whereas for individual control subjects we excluded the subjects’ own data to compute the reference vector. A cosine closer to 1 indicates that the subject-specific and ‘reference’ vectors are more aligned and therefore, the structures of the muscle patterns that they represent are similar.

### Statistical analyses

#### Modulation of muscle activity within groups

Modulation of muscle activity was first evaluated for each group individually. Specifically, we compared muscle activity between the epochs of interest using a Wilcoxon signed-rank test (non-parametric equivalent of paired t-test) for each individual muscle and for each gait cycle phase, resulting in 360 comparisons per epoch (12 intervals × 15 muscles × 2 legs. We subsequently corrected the significance threshold for each epoch using a Benjamini-Hochberg procedure (Benjamini and Hochberg, 1995) to indicate significant changes in our figures, but all data in both groups was used in the structural analyses.

#### Structure of muscle activity patterns during steady state walking

We used a Wilcoxon ranksum test to compare the groups on their ΔEMG_SS_ for each leg during late adaptation in the split-belt condition. We specifically compared the group’s similarity in ΔEMG_SS_ to the reference pattern obtained with the cosine analysis.

#### Sensorimotor recalibration of corrective muscle responses

We compared the regressor coefficients β_no-adapt_ and β_adapt_ for each group to determine if stroke survivors and controls differed in the adaptation of corrective responses. Since the regressor estimates of β_no-adapt_ and β_adapt_ in a regression model are not independent, between-group comparisons were performed in the 2D space covered by β_no-adapt_ and β_adapt_. The differences between the groups were compared using a chi-squared distribution, which could be considered as a high-dimensional t-test (Härdle and Simar, 2007).

#### Correlation analyses

We asked whether individual subjects’ adaptation of muscle activity was related to the severity of motor impairment (i.e. Fugl-Meyer score). To this end, we performed Spearman correlations between 1) the Fugl-Meyer score and 2) outcome measures that reflected sensorimotor recalibration (i.e. β_adapt_ and β_no-adapt_) and the similarity metric comparing the structure of muscle activity during late adaptation in the split-belt condition for each individual vs. a reference ΔEMG_SS_.

#### Modulation of kinematic parameters

We compared stroke survivors and controls in how they modulated kinematic parameters. To this end, we performed a repeated measures ANOVA for each kinematic outcome (StepAsym, StepPosition, StepTime and StepVelocity) with GROUP (stroke vs. Controls), EPOCH (early Adaptation, late Adaptation and early Post-Adaptation) and the interaction between both variables as predictors. Note that we did this analysis with unbiased data (i.e., baseline subtracted) because we were interested in differences in modulation across groups, beyond their baseline biases. In case of a significant GROUP or GROUPxEPOCH interaction effect, we performed between group comparisons for each epoch using Bonferroni corrected independent t-tests (adjusted α=0.017).

#### Speed-matched analysis

Stroke survivors walked slower than controls during the experiment (averaged medium speed= 0.78±0.24 vs. 1.07±0.12 m/s, ranksum test p<0.01). Thus, we repeated our analyses with only the 10 fastest participants in the stroke group and the 10 slowest controls to determine if structural differences between our groups were due to walking speed, rather than brain lesion. Walking speed was not significantly different for these speed-matched subgroups (0.88±0.18 vs .1.0±0.15 m/s, ranksum test p=0.10). Importantly, selection of the fastest stroke survivors did not result in a selection of patients with less severe motor impairments (Fugl Meyer score = 29.5±3.4 vs 28.5±5.1, p=0.67, for subgroup included vs. subgroup excluded in the speed-matched comparison respectively).

## Results

### Cerebral lesions interfered with the structure of muscle activity in a novel walking environment

We computed a similarity metric ΔEMG_SS_, which indicated the similarity between the structure of individual’s muscle activity modulation in steady state walking relative the average pattern in controls (‘reference pattern’). We found that the non-paretic’ leg activity at steady state was similar to the one of controls, whereas the paretic leg was not (Figure 3A). Differences in the structure of muscle activity modulation between the groups can be appreciated in Figure 3B. Specifically, similarity metric ΔEMG_SS_ was lower in the paretic leg compared to controls (Figure 3A; p=0.001) and between-group differences were trending (p=0.057) when comparing the non-paretic leg activity to that of controls. These between-group differences were not observed when patients and controls walked at similar speeds (median ± interquartile range in controls vs. stroke survivors for the non-paretic leg: 0.58±0.26 vs. 0.51±0.22, p=0.47; paretic leg: 0.39+0.13 vs. 0.28±0.18, p=0.1). Interestingly, a more atypical structure in muscle activity modulation in the paretic leg was associated with poorer voluntary leg motor control as measured by the Fugl-Meyer scale (rho = 0.59, p=0.028, Figure 3C), but not in the non-paretic leg (rho = −0.29, p=0.32 data not shown). In conclusion, the structure of muscle activity at steady state was different between patients and controls and individuals with more atypical paretic activity were those with lower voluntary function.

**Figure 3.**
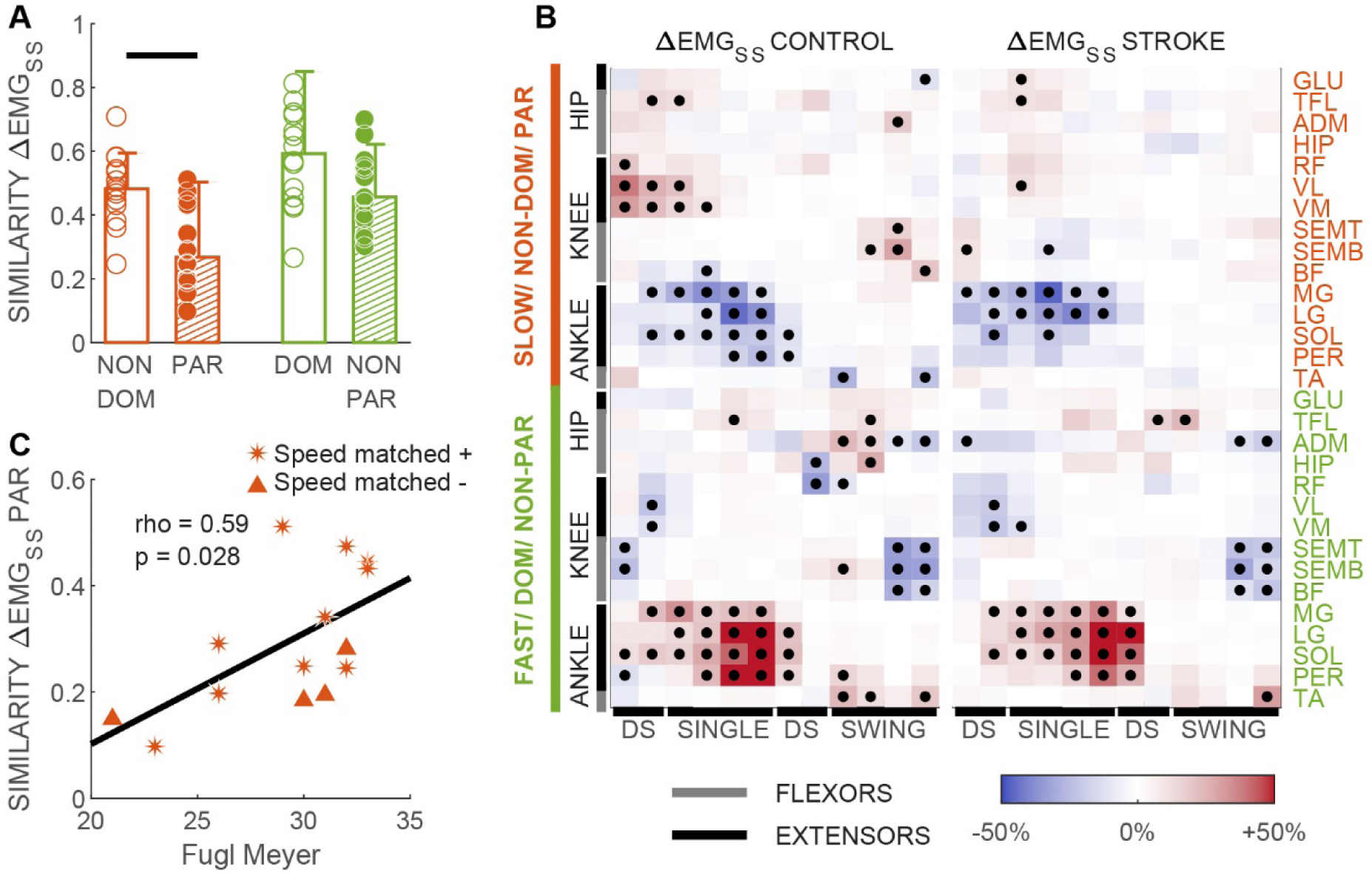
Structure of muscle activity modulation. **A)** Similarity of individual subjects’ steady state muscle activity modulation to the reference pattern (i.e. expressed as the cosine between individual subject vector and group median of controls). Values closer to 1 indicate more similarity between vectors. Bars indicate group medians, error bars represent the interquartile range. Horizontal lines indicate significant differences in group medians as determined with a Wilcoxon Ranksum test (p<0.05). **B)** Visual representation of the structure of muscle activity modulation in the steady state of split-belt walking (ΔEMG_SS_) relative to baseline walking. Red colors indicate increased activity and blue colors indicate decreased activity. Dots indicate statistical significance for non-parametric within group comparisons. We corrected the significance threshold for each epoch using a Benjamini-Hochberg procedure (Benjamini and Hochberg, 1995), setting the acceptable false discovery rate to 10%. In addition, we focused on significant differences between epochs that exceeded 10% of the maximum baseline activity for a given muscle since we considered these to be meaningful changes. Corrected p-thresholds for ΔEMG_SS_ were 0.058 for controls and 0.02 for stroke. **C)** Association between severity of motor symptoms (Fugl Meyer test) and structure of EMG modulation in steady state walking (ΔEMG_SS_). Asterisks represent subjects included in the speed-matched analysis whereas triangles indicated subjects that were excluded. We found a significant correlation (i.e. Spearman’s rho) with less severely affected stroke survivors exhibiting muscle activity modulation closer to the reference pattern.

### Sensorimotor recalibration of corrective responses was intact after cerebral lesion

The structure of corrective responses for each group indicated that on average both groups recalibrated their gait similarly. This is qualitatively indicated by the “checker boards” illustrated in Figure 4. Notice that in both groups the observed corrective responses post-adaptation (Figure 4C) look more similar to those predicted by the adaptive (Figure 4B) than the environment-based modulation (Figure 4A). The environment-based and adaptive-based contributions to corrective responses post-adaptation were quantified with a regression model, which reproduced the data well (Figure 5 left panels). We observed that the regression coefficient β_adapt_ was greater than β_no-adapt_ in both groups for the leg that walked slow (i.e., non-dominant leg in controls: CI for β_adapt_=[0.68-0.85] vs. CI for β_no-adapt_=[0.18-0.35]; paretic leg in stroke: CI for β_adapt_=[0.55-0.77] vs. CI for β_no-adapt_=[0.10-0.32]) and the leg that walked fast (i.e., dominant leg in controls: CI for β_adapt_=[0.73-0.89] vs. CI for β_no-adapt_=[0.09-0.25], non-paretic leg in stroke: CI for β_adapt_=[0.54-0.71] vs. CI for β_no-adapt_=[0.46-0.62]). These coefficients were not different between groups when estimated from the averaged paretic leg activity across stroke survivors vs. that of the non-dominant leg across controls (Chi^2^=3.2, p=0.20), indicating that averaged responses in the slow leg were adapted to the same extent in stroke survivors and controls. Conversely, we found between-group differences when comparing the coefficients of the averaged non-paretic activity in the stroke group vs. that of the dominant leg in the control group (Chi^2^=48.9, p=2.4*10^−11^, Figure 5A, bottom panel). Thus, we observed between-group differences in the regression coefficients for the non-paretic, but not the paretic, compared to control legs.

**Figure 4.**
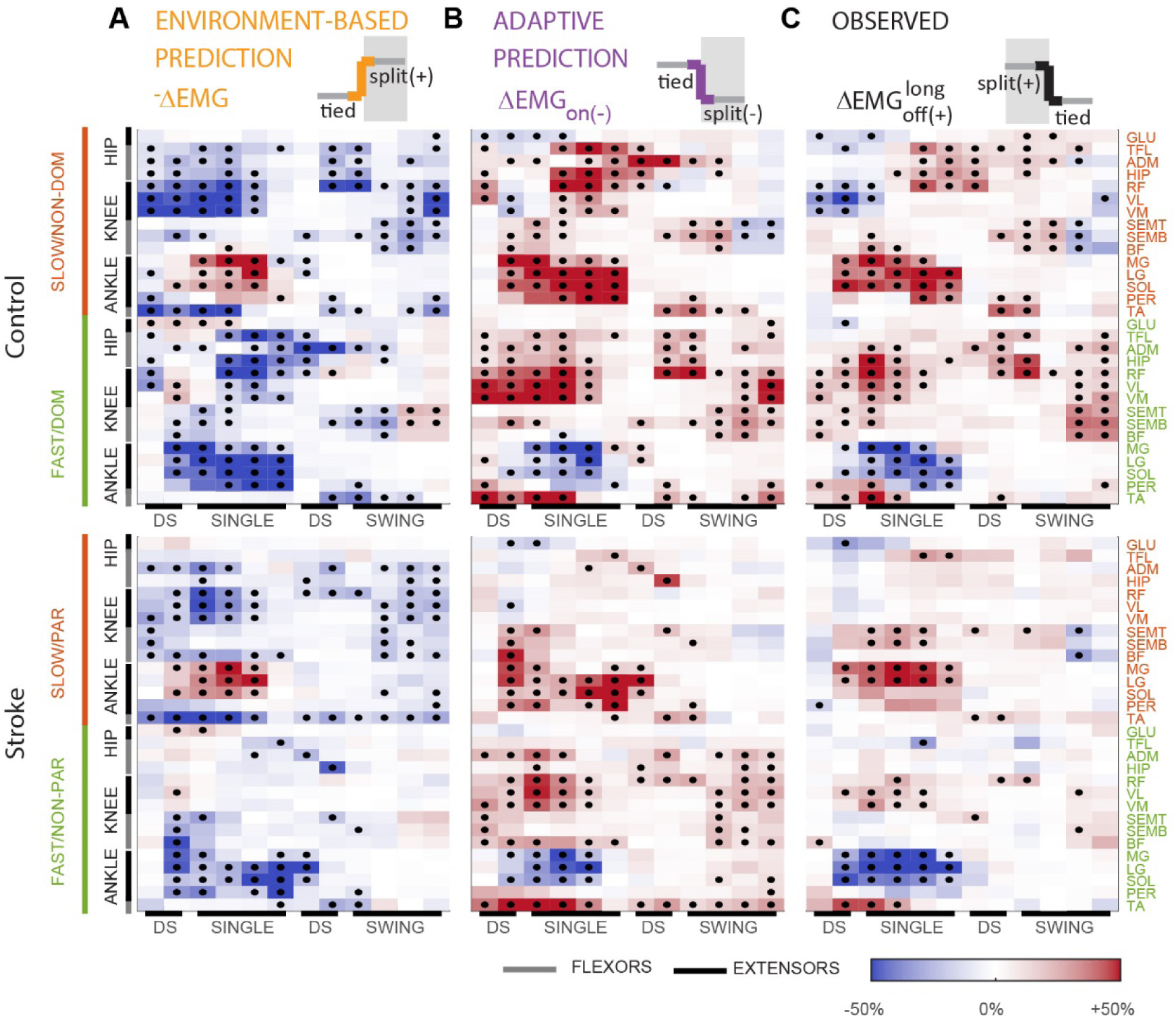
Predicted and measured structure of corrective responses after a long adaptation period. Data of controls are shown in the upper panels and those of the stroke participants in the lower panels. **A**,**B)** Expected corrective responses elicited by the ‘off’ transition under the environment-based (panel A) and adaptive (panel B) cases. Data (in color) and significance (black dots) were derived from the observed corrective responses upon the introduction of the ‘(+)’ walking environment (Supplementary Figure 1), by either taking the numerical opposite (environment-based) or by transposing leg activity (adaptation-based). **C)** Measured corrective responses upon removal of the ‘(+)’ environment.

**Figure 5.**
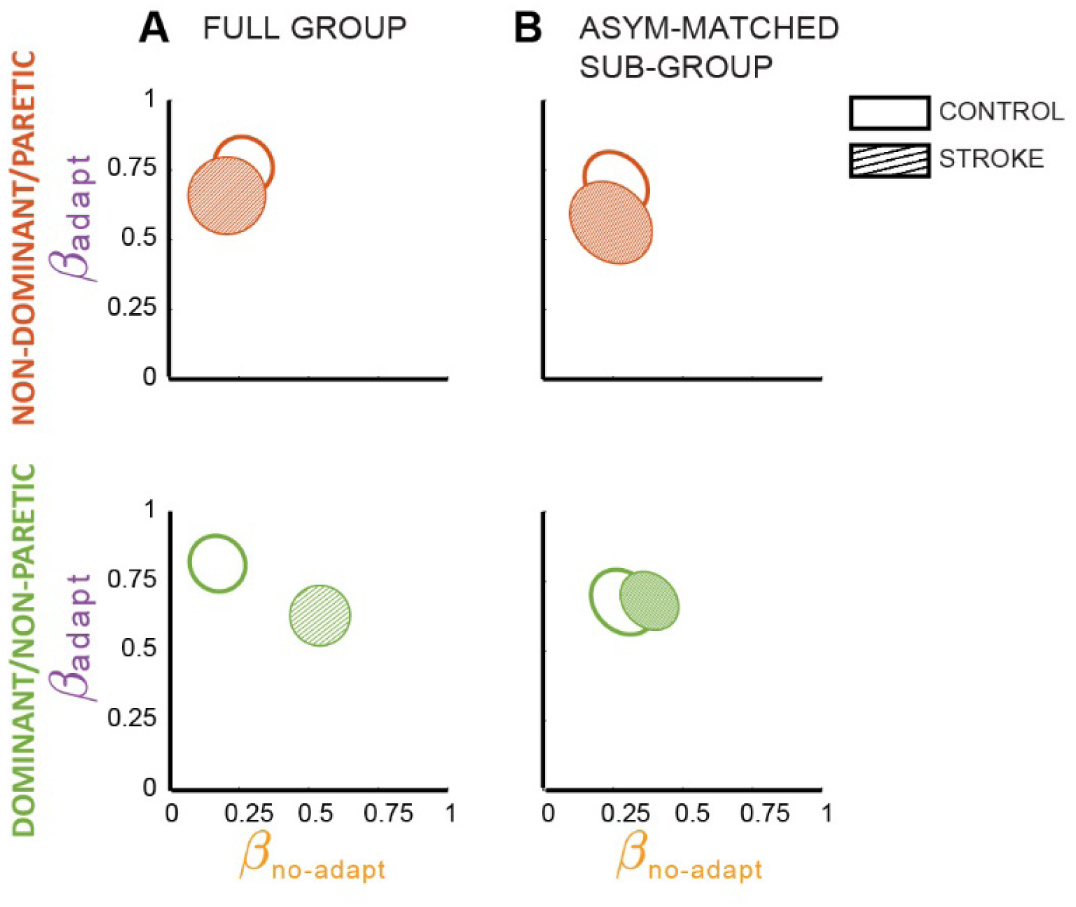
Adaptive and environment-based contributions to corrective responses. The ellipses represent the regression estimations of β_adapt_ and β_no-adapt_ and their 95% confidence intervals for the control group (open) and the stroke group (hatched). **A)** Data obtained with 14 subjects per group. Paretic leg: R^2^= 0.47, model p-value = 8.8*10^−26^. Non-paretic leg: R^2^= 0.67, model p-value = 1.3*10^−45^. Dominant leg in controls R^2^= 0.71, model p-value = 1.1*10^−48^. Non-dominant leg in controls R2= 0.68, model p-value = 3.1*10^−45^. **B)** Data obtained for asymmetry matched groups (i.e. n=7 per group). Non-dominant/paretic leg: controls: CI for β_adapt_ = [0.61 0.79] and CI for β_no-dapt_ = [0.17-0.25], R^2^= 0.64, model p-value = 1.2*10^−40^; stroke: CI for β_adapt_ = [0.45-0.68] and CI for β_no-adapt_ = [0.13-0.36], R^2^= 0.44, model p-value = 6.2*10^−23^); between-group comparison: Chi^2^=4.1, p=0.13. Dominant/ non-paretic leg: controls: CI for β_adapt_ = [0.58 0.77] and CI for β_no-dapt_ = [0.20-0.38], R^2^= 0.63, model p-value = 8.4*10^−39^; stroke: CI for β_adapt_ = [0.60-0.76] and CI for β_no-adapt_ = [0.30-0.46], R^2^= 0.73, model p-value = 8.8*10^−51^); between-group comparison: Chi^2^=2.5, p=0.29.

As a post-hoc analysis, we considered the possibility that these group differences in the non-paretic side could arise from our estimation of the adaptive-based modulation (ΔEMG_on(-)_). Notably, this muscle activity was not recorded but it was inferred from the muscle activity of the other leg, assuming symmetry of corrective responses across legs. Given that stroke survivors exhibit asymmetric motor patterns, the paretic leg’s ΔEMG_on(+)_ may not be a good estimate for the non-paretic leg’s ΔEMG_on(-)_, thereby leading to underestimation of β_adapt_ in this leg. Thus, we performed a subgroup analysis in which stroke survivors and controls were matched for symmetry in their muscle activity during baseline walking. We did not find between-group differences for either leg of the stroke group compared to the controls when asymmetry in baseline muscle activity was matched between the groups (Figure 5B; paretic vs. non-dominant control leg Chi^2^=4.1, p=0.13 and non-paretic vs. dominant control legs Chi^2^=2.5, p=0.29). In conclusion, the observed structure of corrective responses post-adaptation were more similar to the one predicted by adaptive, rather than environment-based, modulation in patients with cerebral lesions and controls.

While recalibration of corrective responses post-stroke did not differ from controls at the group level when asymmetries were accounted for, we considered the possibility that some individuals would exhibit less recalibration compared to others. Consistently, Figure 6A shows a wide range of β_adapt_ and β_non-adapt_ regression values at the individual level. Also, note that the regression model had smaller R^2^ when applied to each subject’s corrective responses post-adaptation (controls’ non-dominant leg: R^2^=0.38±0.18; controls’ dominant leg: 34±0.17; paretic leg: 0.18±18; non-paretic leg: 0.18±18) than to the group’s corrective response (reported in previous section). However, the regression model was significant in all individuals, except for one stroke survivor (p=0.19). In sum, we find large ranges of regression coefficients in control and post-stroke individuals.

**Figure 6.**
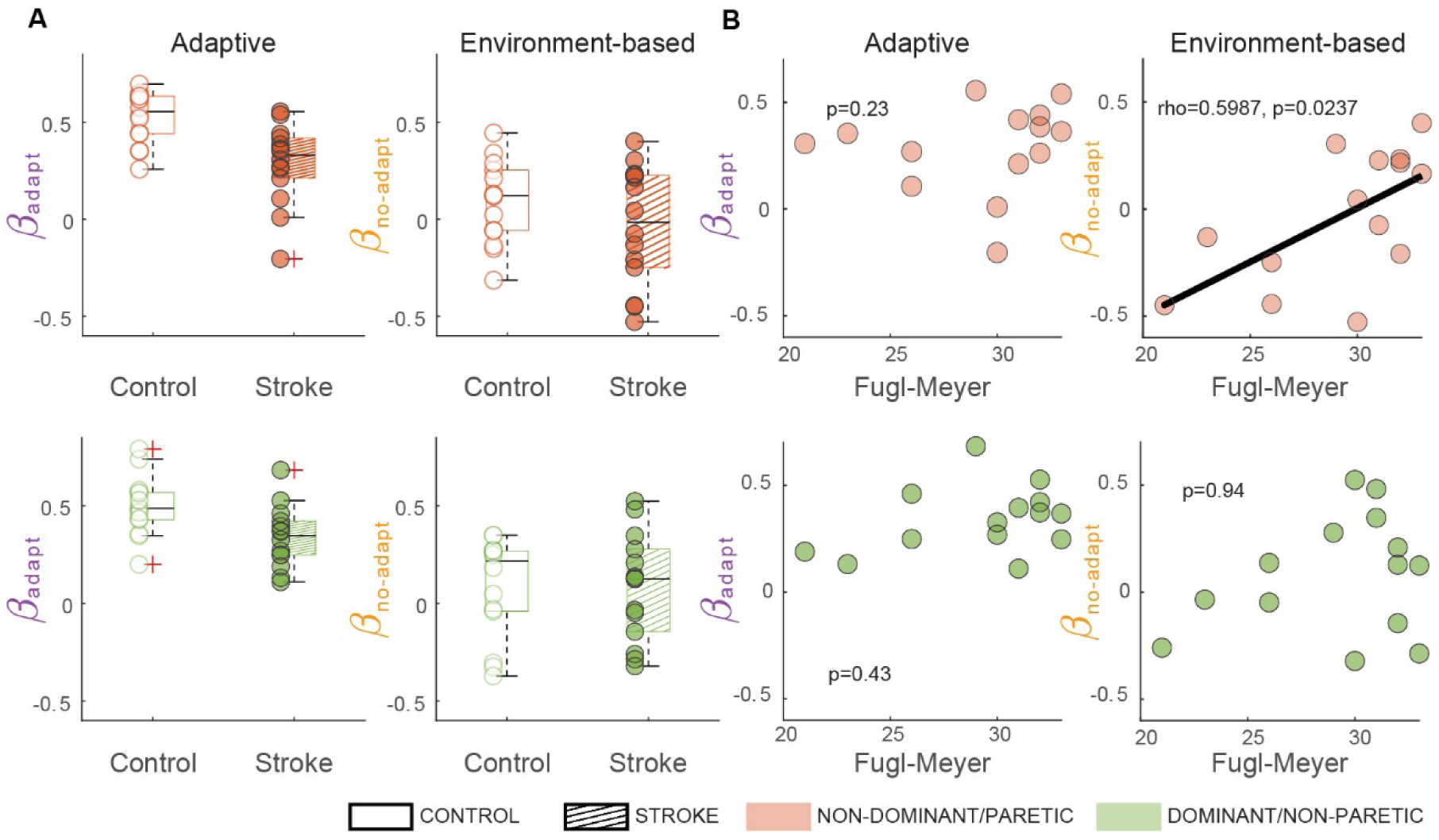
Individual regression results. **A)** Intersubject variability for the adaptive (β_adapt_) and environment-based (β_no-adapt_) contributions to corrective responses in the slow/paretic leg (upper panels) and the fast/nonparetic leg (lower panels). Median ± interquartile range for regressors are as follows: Non-dominant leg: β_adapt_=0.55±0.19; β_non-adapt_=0.12±0.30; p=9.2*10^−19^±7.7*10^−13^. Dominant leg: β_adapt_=0.49±0.14; β_non-adapt_=0.22±0.31; p=1.2*10^−15^±4.7*10^−12^. Paretic leg: β_adapt_=0.33±0.21; β_non-adapt_= −0.02±0.4;p=2.1*10^−8^±3.2*10^−5^. Non-paretic leg: β_adapt_=0.35±0.17; β_non-adapt_=0.13±0.42; p=1.6*10^−9^±6.2*10^−6^. **B)** Spearman correlations between leg motor function (Fugl-Meyer scale) and β_adapt_ and β_no-adapt_ for each leg.

We further asked if stroke survivors would exhibit less recalibration if they had more severe leg motor impairments (i.e. Fugl-Meyer Scale). Thus, we computed the Spearman correlation between individual subjects’ regressors and their leg motor score (Figure 6B). We found that β_adapt_ of neither the paretic or non-paretic legs was correlated to the Fugl-Meyer score (paretic: rho=0.34, p=23; non-paretic: rho=0.23, p=0.43). On the other hand, motor function measured with the Fugl-Meyer score was associated with the paretic’s β_no-adapt_ and not the non-paretic’s β_no-adapt_ (Paretic: rho=0.60, p=0.024; non-paretic: rho=0.02, p=0.94). However, this correlation was driven by the individual with the largest negative β_no-adapt_ since the correlation was no longer significant when this subject was excluded (rho=0.37, p=0.22). As such, we are cautious about interpreting this result as a positive association between environment-based corrective response and leg motor scores. Together our correlation analyses indicate that recalibration of corrective responses is not associated with the quality of voluntary motor control.

### Stroke-related deficits in muscle coordination are not reflected in asymmetry parameters

While stroke survivors exhibited deficits in the execution of updated motor commands during steady state split-belt walking (i.e. ΔEMG_SS_), we observed no differences between the groups in the modulation of asymmetry parameters (i.e. stepAsym, stepPosition, stepTime and stepVelocity, Figure 7). Specifically, we observed no main effects of GROUP or GROUPxEPOCH interaction effects for the interlimb kinematic parameters (p>0.05). Comparable results were obtained in our speed-matched analysis. Thus, interlimb kinematic parameters are less sensitive to stroke-related deficits in locomotor adaptation than our outcome measures for muscle coordination.

**Figure 7.**
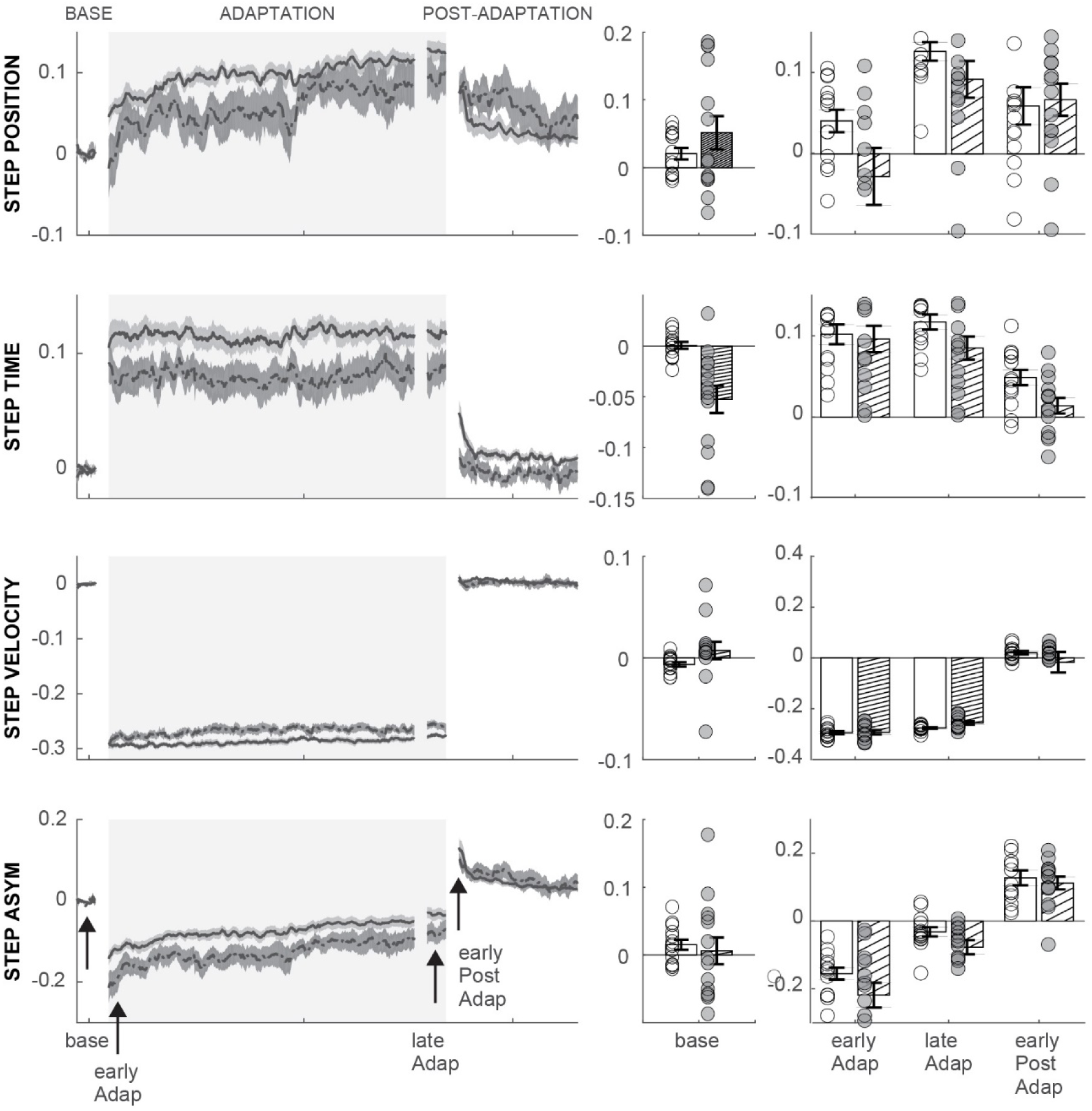
Modulation of kinematic parameters. **A)** Group averaged time courses for StepPosition, StepTime, StepVelocity and StepAsym. Note that individual subjects’ baseline biases were subtracted to allow for comparison of modulation of parameters regardless of differences in baseline asymmetry. Shaded areas represent standard errors for each group. For visual purposes, data were smoothed using a running average (median) of 10 strides. Rectangles represent the epochs of interest. **B)** Interlimb kinematic parameters for each group during baseline. **C)** Between-group comparisons for kinematic parameters over the epochs of interest. We found no significant differences between the groups in any of the parameters. StepAsym (GROUP: F_(1,28)_=3.48, p=0.07; GROUPxEPOCH: F_(2,56.)_=1.16, p=0.31), stepPosition (GROUP: F_(1,28)_=2.14, p=0.16; GROUPxEPOCH: F_(2,56.)_=2.33, p=0.13), stepTime (GROUP: F_(1,28)_=3.17, p=0.09; GROUPxEPOCH: F_(2,56)_=0.63, p=0.50) and stepVelocity (GROUP: F_(1,28)_=1.41, p=0.25; GROUPxEPOCH: F_(2,56)_=1.11, p=0.34).

## Discussion

We studied the involvement of cerebral structures in the sensorimotor recalibration of gait using stroke as a clinical model. We found that on average stroke survivors had intact recalibration of corrective responses in the paretic leg, which was surprising given then known deficits in paretic responses post-stroke. On the other hand, we found cerebral lesions affected the paretic legs’ muscle activity in the steady-state of split-belt walking. Thus, our results suggest that intact cerebral are needed for the execution, but not the recalibration, of motor commands upon novel movement demands.

### Sensorimotor recalibration of corrective responses is intact after cerebral lesions

We found that the adaptive and environment-based contributions to corrective responses were comparable between stroke survivors and controls in the paretic, but not in the non-paretic leg which exhibited lower β_adapt_ estimates. Rather than reflecting a poor recalibration in the non-paretic leg, we speculate that β_adapt_ was underestimated due to asymmetry in corrective responses. Indeed, regression coefficients for either leg were similar across groups with asymmetry-matched participants. While it is possible that motor asymmetry post-stroke impairs the recalibration of corrective responses, we believe that motor asymmetry simply altered the estimation of β_adapt_. Notably, β_adapt_ quantifies the similarity between corrective responses post-adaptation and the expected adaptive-based modulation (e.g. ΔEMG_on(-)_) counteracting the opposite perturbation to the one originally experienced (Iturralde and Torres-Oviedo, 2019). We inferred ΔEMG_on(-)_ for each leg from the corrective responses of the contralateral leg during early adaptation (ΔEMG_on(+)_), exploiting the symmetry of corrective responses between legs. While this approach is adequate for unimpaired individuals (Iturralde and Torres-Oviedo, 2019), asymmetries in corrective responses post-stroke (Marigold and Eng, 2006; de Kam *et al.*, 2018) result in a poor representation of ΔEMG_on(-)_ (i.e., adaptive-based regressor), and thereby underestimation of β_adapt_ for the most asymmetric individuals. Our results indicate that β_adapt_ was underestimated more in the non-paretic than in the paretic leg. This suggests that corrective responses ΔEMG_on(+)_ in the paretic leg are a poor estimate of ΔEMG_on(-)_ in the non-paretic leg. Perhaps this is due to missing activity patterns in the paretic leg corrective responses (de Kam *et al.*, 2018) that cannot account for activity observed in the non-paretic leg. Thus, group differences of the full groups’ non-paretic vs. control legs are likely due to underestimation of β_adapt_, rather than poor sensorimotor recalibration in the non-paretic leg. Taken together, our results suggest that recalibration of corrective responses is not affected in individuals with cerebral lesions, but future studies are needed to determine the potential impact of motor asymmetry on sensorimotor recalibration.

Alternatively, it is possible that discrepancies between the paretic and non-paretic extent of adaptive-based changes reflect leg-specific recalibration. This is supported by the independent recalibration of the legs in hybrid walking (i.e. one leg moving forward faster than the other leg moving backward (Choi and Bastian, 2007)). However, leg-specific adaptation in hybrid walking may result from the peculiar nature of this task and may, therefore, not apply to other locomotor adaptation paradigms. In fact, more recent studies have demonstrated interlimb transfer of adapted motor patterns during split-belt walking forwards with both legs (Krishnan *et al.*, 2017, 2018), which argues against leg-specific recalibration. Moreover, we have observed that subjects adopt a steady state pattern in the split condition that is influenced by ipsilateral and contralateral speed-demands (Iturralde and Torres-Oviedo, 2019), further arguing against leg-specific recalibration. Hence, it is unlikely that recalibration can be affected in only one leg of individuals post-stroke.

We also found that environment-based and adaptive contributions to corrective responses showed a wide range of values across individuals in both the stroke and the control group, which is consistent with previous reports (Iturralde and Torres-Oviedo, 2019). In addition, the regression R^2^ for individual analyses were lower compared to the group averages likely due to the greater noise in corrective responses at an individual level. Consistently, R^2^ increases when more strides are averaged to characterize corrective responses at every transition (Iturralde and Torres-Oviedo, 2019). Our analysis of individual regression coefficients indicated that the severity of motor impairments (i.e., Fugl-Meyer scores) were not associated with smaller adaptive-based modulation of corrective responses. On the other hand, we found that stroke survivors with more severe motor impairments had negative β_no-adapt_ values, which were also observed in controls. These negative values may reflect a generic response to an environmental transition, regardless of its direction. Such a generalized response may consist of a stiffening reaction upon a sudden unexpected environmental transition, similar as the startling-like responses (Oude Nijhuis *et al.*, 2010). In sum, both groups showed comparable inter-subject differences in their ability to adapt corrective responses.

### Cerebral lesions affect the execution of updated motor commands in a new walking environment

We found that stroke survivors exhibited impaired modulation of steady state muscle activity, which was particularly reflected in muscle activity modulation in their paretic leg. Interestingly, aberrant patterns of muscle activity did not impact the modulation of kinematic asymmetry parameters (Reisman *et al.*, 2007), indicating that muscle activity is more sensitive to stroke-related deficits in motor output than kinematic parameters characterizing their asymmetries. Impaired modulation of steady state muscle activity may reflect subjects’ impaired execution of updated motor commands due the lack of leg motor selectivity. Indeed, more atypical steady state muscle activity was associated with poorer leg motor function. Moreover, prior studies also demonstrated that stroke survivors’ muscle activity during steady state walking is impacted by a lack of selective muscle control (Clark *et al.*, 2010). Thus, impaired motor selectivity probably contributes to the aberrant patterns of muscle activity during steady state split-belt walking. Alternatively, the atypical muscle activity patterns in stroke survivors may have resulted from a lower walking speed. Indeed, steady-state muscle activity became more similar across groups in our speed-matched analysis. However, between-group differences in our similarity metric were still substantial after controlling for speed (0.39+0.13 vs. 0.28±0.18, p=0.1) despite a lack of statistical significance. Lastly, the lack of modulation of steady state muscle activity did presumably not result of muscle atrophy, given that muscle groups that lacked modulation in the steady state (i.e. knee extensors) were highly modulated during corrective responses (See Figure S1). We speculate steady state muscle activity depends on neural circuits involved in voluntary motor control, whereas this is not the case for corrective responses (de Kam *et al.*, 2018). Taken together, our results suggest that stroke survivors exhibit impaired execution of updated motor commands in the steady state of split-belt walking, most likely due to their impaired motor function.

### Partial dissociation between recalibration and execution of updated motor commands

We found that stroke survivors exhibited intact recalibration of corrective responses, but impaired muscle patterns at steady state split-belt walking, suggesting partial dissociation between motor performance in the altered environment and post-adaptation behavior. This finding is consistent with previous work demonstrating that the extent to which subjects adapt their movements during split-belt walking does not predict their after-effects (Sombric *et al.*, 2019). Partial dissociation between steady-state and post-adaptation behavior is further supported by the findings that after-effects are not sensitive to manipulation of steady-state behavior through visual feedback (Wu *et al.*, 2014; Long *et al.*, 2016). Taken together, our findings suggest that steady-state and post-adaptation behaviors are partially independent, and possibly mediated through distinct neural processes.

We found that post-adaptation muscle activity was indicative of sensorimotor recalibration of corrective responses also observed in previous studies (Maeda *et al.*, 2018; Iturralde and Torres-Oviedo, 2019). Since recalibration has also been observed in feedforward motor commands upon perturbation removal (Tseng *et al.*, 2007; Taylor and Ivry, 2014), our results provide further evidence for shared internal models for generating corrective responses and feedforward motor commands (Wagner and Smith, 2008; Yousif and Diedrichsen, 2012; Cluff and Scott, 2013; Maeda *et al.*, 2018). Feedforward adaptation and learning of internal models require intact cerebellar function (Martin *et al.*, 1996; Smith and Shadmehr, 2005; Morton and Bastian, 2006). Most likely, corrective responses depend on spinal cord and brainstem circuits for their execution (Bolton, 2015; Weiler *et al.*, 2019) and on the cerebellum for their adaptation, which would explain why our participants with cerebral lesions showed intact recalibration of corrective responses.

Our observation of stroke-related impairments in steady-state movement execution suggest that these processes are cerebral-dependent, perhaps through connections between cerebral and cerebellar structures (Kelly and Strick, 2003; Hoshi *et al.*, 2005). Moreover, intact motor pathways for voluntary motor control (e.g. corticospinal tract) are most likely involved in the execution of steady-state motor commands (Schweighofer *et al.*, 2018), given our finding that individuals with poorer voluntary motor control also exhibited a more atypical structure of their steady state muscle activity. Such associations were not found for the execution of corrective responses (de Kam *et al.*, 2018), suggesting that the execution of corrective responses uses different circuitry, most likely at the level of the brainstem (Jacobs and Horak, 2007; Bolton, 2015).

Taken together our results are consistent with the idea that corrective and planned actions share an internal model, which relies on cerebellar structures for their adaptation and on cerebral structures for their execution.

## Limitations

Our study has a few limitations. First, because of the lower preferred walking speed in stroke survivors, our participants with stroke performed the experiment at slower walking speed then controls. To assess the confounding effect of velocity, we performed an additional subgroup analysis in which stroke survivors and controls were more comparable in walking speed. In this analysis, we still observed a trend towards impaired execution of updated motor commands in the paretic leg of stroke survivors. We therefore believe that the execution of updated motor commands in steady-state walking is a stroke-related deficit and not merely an effect of slower walking. A second limitation was that ΔEMG_on(-)_ was not directly measured. Instead, ΔEMG_on(-)_ was estimated from ΔEMG_on(+)_ in the contralateral leg. Consequently, any asymmetry in muscle activity would lead to a bad predictor of the corrective responses that would result from recalibration (i.e., EMG_on(-)_) and thereby reduce the possible β_adapt_. Our asymmetry-matched analysis showed that the regression results were indeed influenced by stroke survivors’ asymmetry. Therefore, it is recommended to measure corrective responses to both the ‘+’ and ‘-’ transitions in future studies.

### Clinical implications

Our results have important clinical implications for stroke survivors. First, our detailed characterization of muscle activity modulation during and after split-belt walking allows for the identification of muscle activity that could potentially be targeted by split-belt treadmill training. Moreover, we have shown that the extent to which movements are recalibrated varied greatly across post-stroke individuals. We speculate that individual differences in sensorimotor recalibration may explain why some stroke survivors improve their gait symmetry in response to repeated split-belt treadmill training while others do not (Reisman *et al.*, 2013; Betschart *et al.*, 2018; Lewek *et al.*, 2018). If so, it may be possible to identify patients that will benefit from split-belt training within just a single session. Future studies are needed to determine if individuals’ recalibration of corrective responses can predict their response to repeated training.

---

**Supplementary Table 1.** Muscle abbreviations.

| **Muscle name** | **Abbreviation** |
| --- | --- |
| Tibialis anterior | TA |
| Peroneus longus | PER |
| Medial gastrocnemius | MG |
| Lateral gastrocnemius | LG |
| Soleus | SOL |
| Biceps femoris | BF |
| Semitendinosus | SMT |
| Semimembranosus | SMB |
| Rectus femoris | RF |
| Vastus lateralis | VL |
| Vastus medialis | VM |
| Iliopsoas and sartorius | HIP |
| Adductor magnus | ADM |
| Gluteus medius | GLU |
| Tensor fasciae latae | TFL |

**Figure S1.**
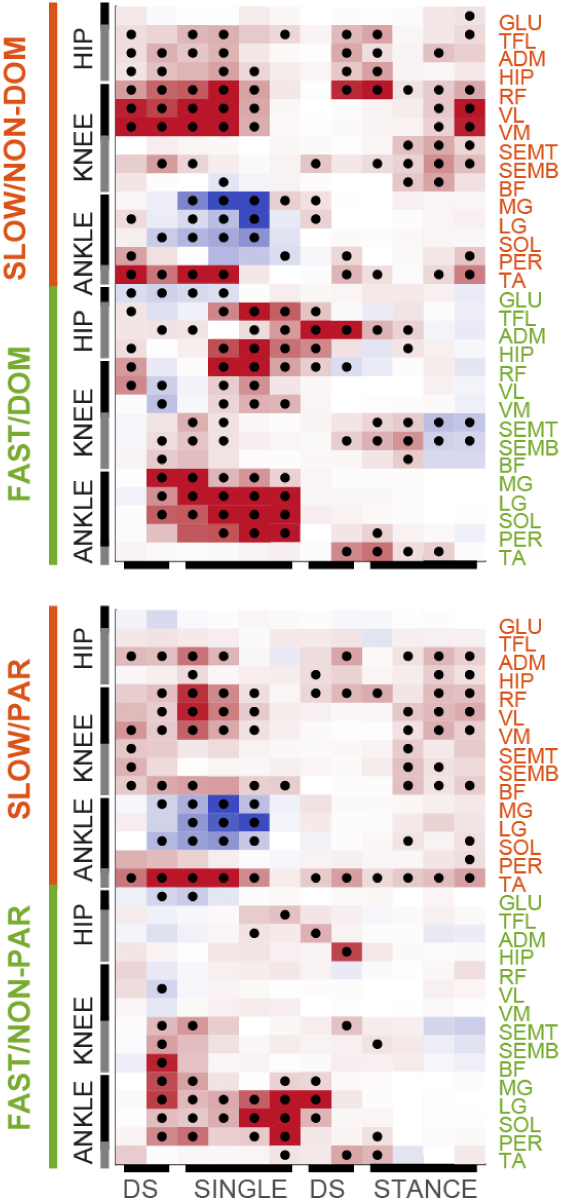
Corrective responses. Corrective responses upon introduction of the ‘(+)’ environment. Upper panel represents controls, lower panel represents stroke survivors. Colors represent group median increase (red) or decrease (blue) in activity. Black dots indicate statistical significance.

